# Mutational and transcriptional landscape of pediatric B-cell precursor lymphoblastic lymphoma

**DOI:** 10.1101/2023.12.27.573433

**Authors:** Emma Kroeze, Ingram Iaccarino, Michelle M Kleisman, Mayukh Mondal, Thomas Beder, Mouhamad Khouja, Marc P Hoeppner, Marijn A Scheijde-Vermeulen, Lennart A Kester, Monika Brüggemann, Claudia D Baldus, Gunnar Cario, Reno S Bladergroen, Nathalie Garnier, Andishe Attarbaschi, Jaime Verdu-Amoros, Rosemary Sutton, Elizabeth MacIntyre, Kenneth Scholten, Laura Arias Padilla, Birgit Burkhardt, Auke Beishuizen, Monique L den Boer, Roland P Kuiper, Jan LC Loeffen, Judith M Boer, Wolfram Klapper

**Affiliations:** Princess Máxima Center for Pediatric Oncology, Utrecht, the Netherlands; Department of Pathology, Hematopathology Section and Lymph Node Registry, University of Kiel, Kiel, Germany; Clinical Research Unit “CATCH ALL” (KFO 5010/1) funded by the Deutsche Forschungsgemeinschaft, Bonn, Germany; Institute of Clinical Molecular Biology, University of Kiel, Kiel, Germany; Centre for Genomics, Evolution & Medicine, Institute of Genomics, University of Tartu, Riia 23b, 51010 Tartu, Tartumaa, Estonia; Medical Department II, Hematology and Oncology, University Hospital Schleswig-Holstein, Kiel, Germany; Department of Pediatrics, Berlin-Frankfurt-Münster ALL Study Group Germany (BFM-G), University Medical Center Schleswig-Holstein, Campus Kiel, Kiel, Germany; Institut d’Hematologie et d’Oncologie Pediatrique, Hospices Civils de Lyon, Lyon, France; Department of Pediatric Hematology and Oncology, St. Anna Children’s Hospital, Medical University of Vienna, Vienna, Austria; Department of Pediatric Hematology and Oncology, Hospital Clínico Universitario de Valencia, Valencia, Spain; INCLIVA, Biomedical Research Institute, Valencia, Spain; Children’s Cancer Institute, University of New South Wales, Sydney, New South Wales, Australia; Laboratory of Onco-Hematology, Necker Children’s Hospital, Assistance Publique-Hôpitaux de Paris (AP-HP), Paris, France; Université Paris Cité, CNRS, INSERM U1151, Institut Necker Enfants Malades (INEM), Paris, France; Paediatric Haematology and Oncology, University Hospital Muenster, Germany; NHL-BFM Study Center, University Hospital Muenster, Germany; Erasmus Medical Center, Sophia Children’s Hospital, Rotterdam, the Netherlands; Department of Genetics, Utrecht University Medical Center, Utrecht University, The Netherlands

## Abstract

Pediatric B-cell precursor (BCP) lymphoblastic malignancies are neoplasms with manifestation either in bone marrow/blood (BCP acute lymphoblastic leukemia, BCP-ALL) or less common in extramedullary tissue (BCP lymphoblastic lymphoma, BCP-LBL). Although both presentations are similar in morphology and immunophenotype molecular studies are virtually restricted to BCP-ALL so far. The lack of molecular studies on BCP-LBL is probably due to its rarity and the restriction to tiny, mostly formalin-fixed paraffin embedded (FFPE) tissues. Here we present the first comprehensive mutational and transcriptional analysis of what we consider the largest BCP-LBL cohort described to date (n=97). Whole exome sequencing indicates a mutational spectrum of BCP-LBL strikingly similar to that found in BCP-ALL. However, epigenetic modifiers were more frequently mutated in BCP-LBL, whereas BCP-ALL was more frequently affected by mutation in genes involved in B-cell development. Integrating copy number alterations, somatic mutations and gene expression by RNA-sequencing revealed virtually all molecular subtypes originally defined in BCP-ALL to be present in BCP-LBL too, with only 7% of lymphomas that were not assigned to a subtype. Therefore, the results here described may pave the way for molecular risk adapted treatment protocols for BCP-LBL patients.

**Keypoints:** Comprehensive molecular characterization of B-cell precursor lymphoblastic lymphoma allows molecular subtyping analogous to leukemias

Compared to leukemias, lymphomas show more alterations in epigenetic modifiers and less in B-cell development genes

## Introduction

B-cell precursor acute lymphoblastic leukemia (BCP-ALL) and lymphoblastic lymphoma (BCP-LBL or B-LBL) are considered one disease entity^1^. Although indistinguishable in morphology and immunophenotype, the clinical presentation is clearly different. BCP-LBL typically manifests as extramedullary disease, localized in the lymph nodes, bones, and skin/subcutaneous tissues. By definition, BCP-LBL patients present with less than 25% of blasts in the bone marrow (BM)^2^. In contrast, BCP-ALL presents as leukemic disease and patients have per definition 25% or more blasts in the BM, often accompanied by peripheral blood involvement and hepato- and/or splenomegaly.^2^

BCP-ALL is one of the most prevalent hematologic malignancy in children^3,4^ and has been extensively studied both clinically and molecularly^5^. Accordingly, treatment strategies for BCP-ALL have evolved by defining molecular genetic subtypes which are associated with outcome, independently of minimal residual disease (MRD)^6^. The BCP-ALL subtype definition is based on aneuploidies, transcription factor (TF)-driven and kinase/cytokine receptor-driven translocations. The high hyperdiploid (HeH) subtype, characterized by 51-67 chromosomes, is one of the most common pediatric ALL subtypes (25%)^6^. Other aneuploid BCP-ALL subtypes include near haploidy (24-30 chromosomes), low hypodiploidy (31-39) and intrachromosomal amplification of chromosome 21 (iAMP21), each accounting for 1-2% of pediatric BCP-ALL. The TF-driven subtypes include *ETV6*::*RUNX1* (25%), *TCF3*::*PBX1* (5-6%), *KMT2A* rearrangements (2%) and also less common subtypes with genetic aberrations of *DUX4*, *MEF2D*, *ZNF384*, *PAX5* and *IKZF1*. Lastly, kinase/cytokine receptor-driven BCP-ALL subtypes include *BCR*::*ABL1* (∼3%), ABL-class (∼3%) and subtypes based on *CRLF2* rearrangements^6^. Frequent secondary genetic events in BCP-ALL include mutations and deletions in genes involved in B-cell development such as *IKZF1* and *PAX5* as well as cell cycle-, RAS- and JAK-STAT pathway genes^7^.

BCP-LBL constitutes a rare hematologic malignancy in children. Currently, patients are stratified based on disease stage as biological risk-factors have not been identified yet^2^. The diagnosis of BCP-LBL is based on mostly small tissue formalin-fixed paraffin embedded (FFPE) biopsies. Although several cytogenetic subtypes have been reported in BCP-LBL, including HeH, *KMT2A* rearrangement, *ETV6*::*RUNX1*, *TCF3*::*PBX1*, and iAMP21^2,8–12^, their incidence and the association with mutations and outcomes remains largely unknown.

## Methods

### Patients’ samples and processing

Pediatric BCP-LBL patients diagnosed between 1996-2022 were included in the study (n=97). Patients’ tumor tissue was obtained from fresh frozen (n=15) or FFPE (n=82) tissue. Clinical data was retrieved from the contributing hospitals. Patient samples were processed at the Princess Máxima Center for Pediatric Oncology, the Netherlands (n=40) and at the Haematopathology Section of the University Hospital Schleswig-Holstein, Germany (n=57). The study was conducted in accordance with the declaration of Helsinki and all patients or patients’ guardians signed informed consent according to local guidelines. Genomic DNA and RNA was extracted from sections of tumor biopsies with at least 40% of tumor cell content as determined by a pathologist using HE staining. Genomic DNA was used for NGS library preparation using either the KAPA Library Preparation kit or the KAPAHyperPlus kit (KAPA Biosystems, MA, USA). Genomic libraries were captured with the HyperPlus Capture Exome Kit or the xGen Exome Panel V2 (Integrated DNA Technology, USA) and sequenced on the Illumina Novaseq6000 (2x150bp paired-end). Some of the libraries (n=52) were also analyzed with the EuroClonality-NDC Assay (Univ8 Genomics. Belfast, UK)^13^. For transcriptome sequencing (RNAseq), RNA libraries were generated using either the KAPA RNA HyperPrep Kit with RiboErase (KAPA Biosystems, MA, USA) and sequenced directly, or with the xGen Broad-Range RNA Library Prep Kit (Integrated DNA Technology, USA) and captured with the xGen Exome Panel V2 (Integrated DNA Technology, USA) to select for coding sequences only. A detailed description of the methodological procedures is provided as **Supplemental Methods**.

### Data analysis

For single nucleotides variants analysis, somatic mutations were called using Mutect2 from GATK 4.2.0.0^14^. Due to the absence of germline material, a selection of 536 genes including BCP-ALL candidate genes^7^, genes with a known function in DNA repair and genes frequently mutated in lymphomas^15,16^ were used for variant analysis (**Supplemental Table 1**). Mutations that passed filtering steps and that were found to occur in ≥5% of the samples were used for further analysis. Copy number aberrations were called by implementing GATK v4.0.1.2 best practices^17^. RNAseq data were processed using STAR v2.7.2d, and STAR-Fusion v1.8.0 and/or Arriba were used for gene fusion detection^18,19^. Count matrices were generated by implementing the featurecounts function from the Rsubread v1.6.4 package. Transcript per million (TPM) normalized data was used to predict subtypes using the ALLCatchR classifier^20^. Subtypes were assigned by integrating information coming from aneuploidies, fusions, single nucleotides variants (SNVs) and ALLCatchR prediction. Samples with inconsistencies in data coming from the different sources were defined as BCP-LBL not otherwise specified (NOS). Samples for which lack of experimental data prevented subtype definition were labelled as “n.a.” (not assigned). For those cases with missing experimental data but assigned to a subtype with “high-confidence” by ALLCatchR^20^, the ALLCatchR prediction was used as final subtype.

For immunoglobulin (IG) and T-cell receptor (TR) rearrangement analysis, sequencing data obtained using the EuroClonality-NDC Assay (Univ8 Genomics. Belfast, UK) were analyzed using the ARResT/Interrogate bioinformatic platform (http://arrest.tools/interrogate-latest/).

To compare data obtained in the analysis of our BCP-LBL cohort with similar data obtained on pediatric BCP-ALL, we used as reference a cohort of 2148 patients (aged 1-18 years) published by Brady et al. (2022)^7^. A more detailed description of data analysis is provided as **Supplemental Methods**.

## Results

### Study cohort

We collected a retrospective cohort of 97 pediatric BCP-LBL patients diagnosed between 1996 and 2022. The median age at diagnosis was 7 years (range 1-18) with a male to female ratio of 0.96:1 (51% female). The majority of the patients were treated with ALL-based treatment protocols, consisting of three chemotherapy blocks (induction, consolidation and maintenance). Patients with higher stages of disease (stage III/IV) received an extra treatment block (reinduction). Lumping major groups of tissues analyzed reveals lymph node in 63%, skin/subcutaneous lesions in 26% and bone lesion in 38% of patients. By definition BM involvement was <25%. Our cohort was comparable to previously described BCP-LBL cohorts, based on median age and disease localizations, as well as distribution of stages and outcome (**Supplemental Table 2**)^2,21^. Nine patients had an event, including 7 relapses, 1 death due to therapy complications and one death in remission due to a secondary malignancy. The median time to follow-up was 61 months, and the 5-year event-free survival rate of the patients in our cohort was 88%±4%, again comparable to previously described cohorts (**Supplemental Table 2**)^2,22^. A more detailed description of patient characteristics is provided in **Supplemental Table 3. Figure 1** shows a schematic depiction of the study design, the cohort characteristics and of the NGS analyses performed on the patient samples.

**Figure 1:**
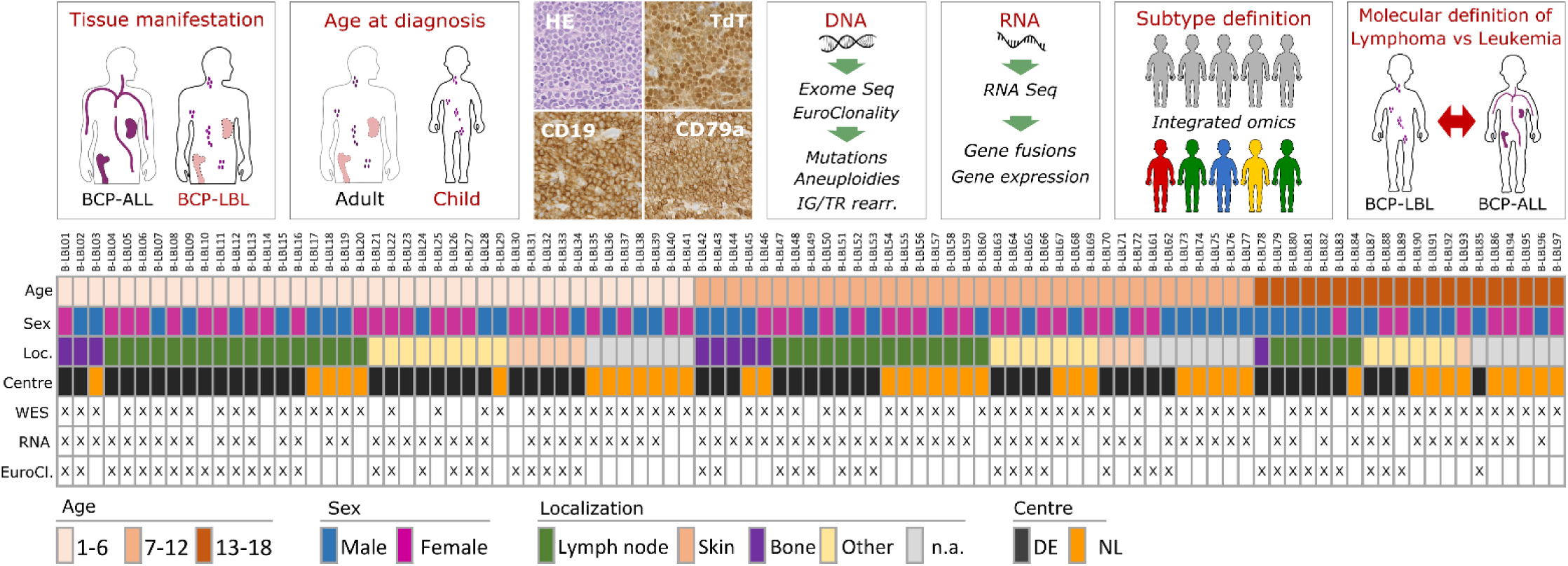
Graphical description of the study design and the experimental cohort. Patients eligible for this study were selected according to the criteria defined in the upper panels of this figure. Selected patients had a degree of bone marrow (BM) involvement lower that 25%; an age at diagnosis between 1 and 18 years old; BCP-LBL cellular morphology, positivity for TdT and CD19 and/or CD79a. DNA and RNA was extracted from tissue biopsies of the selected patients and processed for DNA and RNA next generation sequencing (NGS) analyses. The results of these analyses were integrated to assign lymphoma subtypes and define molecular differences between the BCL-LBL cohort here described and publicly available data obtained from pediatric BCP-ALL patients (Brady et al. 2022). The panel below shows for each of the 97 patients, information about the age group, sex, location (Loc.) of the biopsy used for molecular analyses, tissue processing centre (DE, Germany; NL, Netherlands) and the NGS analysis performed (WES, whole exome sequencing; RNA, whole transcriptome sequencing; EuroCl., targeted sequencing of selected genomic regions using the EuroClonality NDC Assay).

In line with the criteria for BCP-LBL pathology classifications^1,23^, all included patients were positive for B-cell markers at immuno-histological (IHC) inspection (e.g., CD19, CD79a) and all of them, except three, expressed the precursor-marker TdT. Lack of TdT expression by IHC in two out of three samples was presumably rather a technical issue, because expression levels of the TdT gene (DNTT) from RNA from the same samples were in line with the rest of the cohort (data not shown). Furthermore, 6% of the patients showed expression of T-cell marker CD3, and 20% had expression of myeloid marker myeloperoxidase (MPO, **Supplemental Table 3**). The frequency of MPO expression in comparable to a previously published cohort of lymphomas^24^.These specimens were nevertheless included in the study as aberrant expressions are common and the patients were clinically managed as BCP-LBL. As reported in **Supplemental Table 3**, clinical characteristics of 81 patients of our sequencing cohort have been reported in Au-Yeung et al.^9^ (n=44) and Kroeze et al. (n=37)^2^.

### IG and TR rearrangements

B-cell maturation and differentiation is associated with stages of IG rearrangements^25–27^. In early stages of differentiation, recombination of the immunoglobulin heavy chain gene (IGH) gene is either incomplete or unproductive. We therefore sought to use rearrangements analysis as a tool to validate the precursor origin of the BCP-LBL of our cohort, by looking at frequency and productivity of IG/TR rearrangements. IGH and TR gene rearrangements were analyzed in 52 samples (**Figure 1**). As expected for a lymphoid neoplasm, all patients had at least one clonal rearrangement. Most of the patients had rearrangements in the *IGH* and *TRD* genes (89% and 79% respectively). Rearrangements in other IG and TR genes could be identified at different frequencies in our cohort (**Figure 2**). Seventy-eight percent of the total clonal rearrangements identified in the different IG and TR genes in all BCP-LBL samples could be considered either unproductive or non-functional (**Figure 2, Supplemental Table 4**). These results confirm the B-cell precursor origin of BCP-LBL and does not indicate a different stage of maturation of the cell or origin in BCP-LBL compared to BCL-ALL^28^.

**Figure 2:**
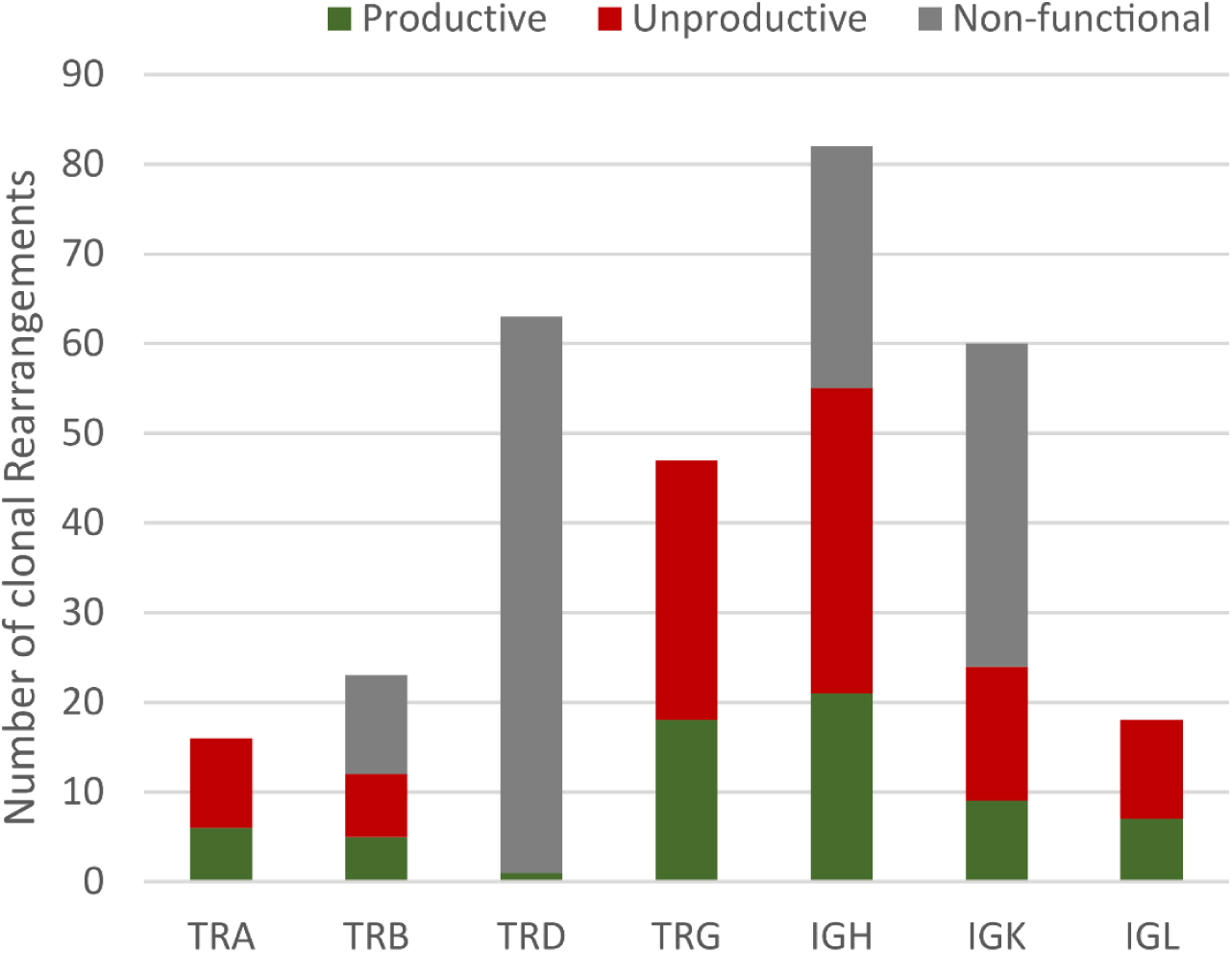
IG/TR Rearrangements analysis. The figure shows the analysis of results obtained using the EuroClonality NDC capture panel on genomic DNA from 52 patients of the BCL-LBL cohort. For each of the T cell receptor genes (TRA, TRB, TRD and TRG) and immunoglobulin genes (IGH, IGK and IGL) the number of clonal rearrangements identified is shown. Furthermore, information from the rearrangement sequence and segment usage was integrated to count the number of rearrangements defined as productive (complete rearrangements, potentially able to produce a functional receptor), unproductive (complete rearrangements unable to produce a receptor for the presence of out of frame deletions) and non-functional (incomplete, chimeric or disruptive rearrangements).

### Mutational landscape and focal deletions in BCP-LBL

Whole exome sequencing (WES) was performed on FFPE material or frozen tissue (n=80). For 13 additional patients, mutational data on a limited set of lymphoma-related genes, covered by the Euroclonality NDC capture assay, was available. The recurrently mutated and deleted genes in BCP-LBL were involved in cell cycle regulation, RAS signaling, B-cell development and epigenetic regulation, and include *CDKN2A/B* (21%), *NRAS* (13%), *IKZF1* (12%), and *KMT2D* (12%) as the most commonly altered genes (**Figure 3**). Next, we asked if our BCP-LBL cohort was characterized by gains or losses of chromosomal arms or complete chromosomes. Analysis of the WES data revealed that aneuploidies were common in our patient cohort, with 83% of the patients showing one or more chromosomal copy number alteration (CNA), including gain of chromosomes typical for HeH and iAMP21 (**Figure 4**). Commonly gained chromosomes included chromosomes 21 (in 38% of the patients), 14 (28%), 6 (23%) and 17 (16%). Chromosome arms affected by recurrent deletions included 9p (11%), 7p (11%) and 13q (6%).

**Figure 3:**
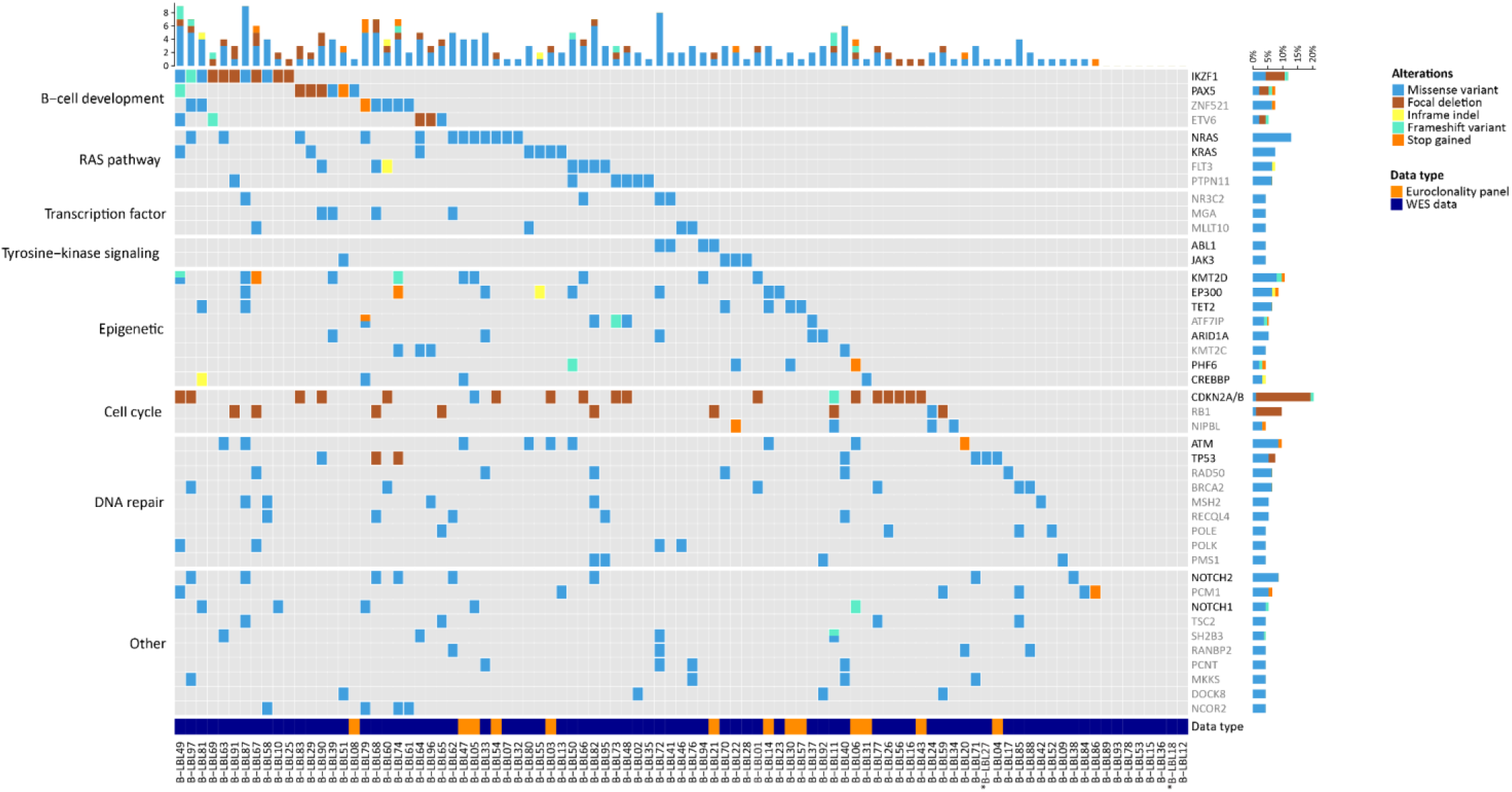
Overview of recurrently mutated genes and pathways in BCP-LBL. Oncoplot showing recurrently mutated and deleted genes in our BCP-LBL cohort. Mutated genes were divided in functional categories as indicated on the left of the figure. For 80 patients, mutational data derives from whole exome sequencing analysis (data type: WES). The analysis of WES data was limited to a selection of 536 genes with known oncogenic potential (see Methods). For 13 additional patients, mutational data derives from the analysis of a limited set of lymphoma-related genes (in black in the gene list), covered by the Euroclonality NDC capture assay (data type: Euroclonality panel). For samples with an asterisk, focal deletions could not be determined.

**Figure 4:**
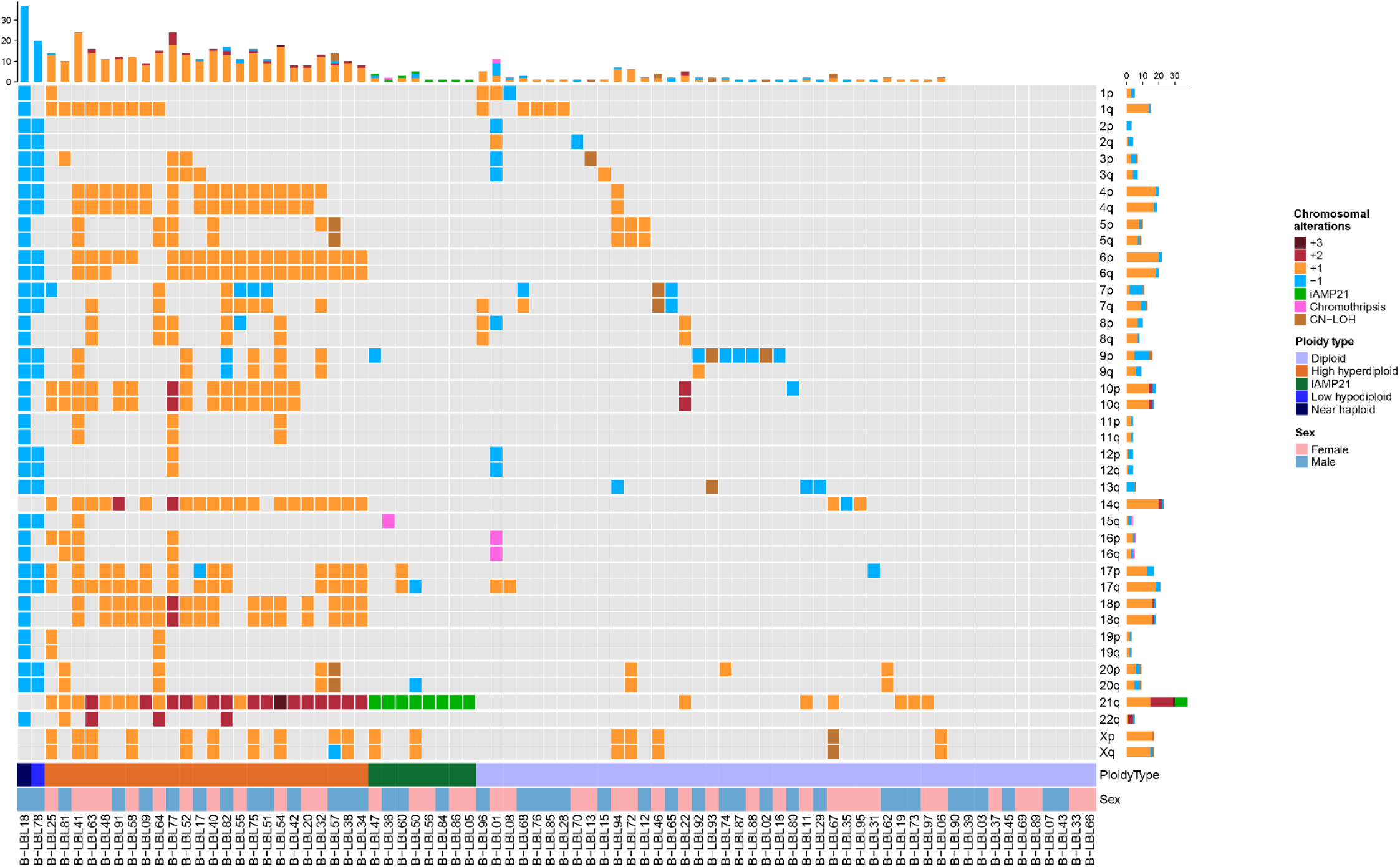
Overview of chromosomal gains and losses in BCP-LBL. Oncoplot showing the chromosomal aberrations per chromosomal arm. Different colors represent number of gains/losses or other chromosomal events.

### Gene fusions

Gene fusions are commonly observed in BCP-ALL and are considered to be driving oncogenic events. To identify chimeric transcripts resulting from gene fusion events, as well as transcriptional signatures defining distinct subgroups, we performed whole transcriptome analysis on RNA extracted from 80 BCP-LBL patients. We found that 45/80 (56%) of the patients carried known or novel gene fusions. The *ETV6*∷*RUNX1* fusion was the most frequent fusion, observed in ten patients, followed by fusions involving *TCF3* (seven patients), with five patients carrying a *TCF3*∷*PBX1* fusion, one patient with a *TCF3*∷*FLI1* fusion and one patient a *TCF3*::*BRD3* fusion. Other TFs involved in fusion genes included *KMT2A* and/or *MLLT* genes (five patients), *NUTM1* (three patients), and *PAX5* (four patients). Rearrangements of *DUX4* to the *IGH* locus were detected in three patients, with three other patients suspected as *DUX4*-rearranged based on high *DUX4* expression and/or a detected breakpoint in the RNAseq reads. Two patients had a *BCR*::*ABL1* fusion and one patient an *ETV6*::*ABL1* fusion. Interestingly, four novel fusions were identified, being *PAX5*::*GLIPR2*, *TCF3*::*BRD3*, *MTO1*::*JPH3*, and *ZNF710*::*CRTC3* (**Supplemental Table 5**; **Supplemental Figure 1**). The patients with *PAX5*::*GLIPR2* and *MTO1*::*JPH3* fusions had a HeH karyotype, for the remaining patients with novel fusions, no other driving lesions were detected.

### Comparison of mutations in BCP-LBL and BCP-ALL

We compared the frequencies of the most frequently mutated genes in our BCP-LBL cohort, to those reported for pediatric BCP-ALL by Brady et al. (2022)^7^. To highlight differences in functional categories, we focused on the top mutated genes of each category defined in **Figure 5 and Supplemental Table 6**. Interestingly, we observed an enrichment in genes with a known role in epigenetic regulation among all genes that were significantly more often mutated in BCP-LBL compared to BCP-ALL. These included *KMT2D* (p=0.0047), *EP300* (p<0.0001), *ARID1A* (p<0.0001) and *ATF7IP* (p<0.0001) among those most frequently mutated, but also *SETD1B* (p=0.0008) and *KMT2C* (p=0.0030) (**Figure 5**). In contrast, TFs important for B-cell development like *PAX5* and *ETV6* were significantly more frequently altered in BCP-ALL (p=0.0011, p=0.0270, respectively) (**Figure 5 and Supplemental Table 6**). Finally, we observed relatively frequent mutations in DNA repair genes, with mutations in *ATM* being significantly more frequent in BCP-LBL compared to BCP-ALL (p<0.0001). Mutations in RAS pathway genes and the tumor suppressor genes *IKZF1*, *RB1* and *TP53* showed similar frequencies in BCP-LBL and BCP-ALL.

**Figure 5:**
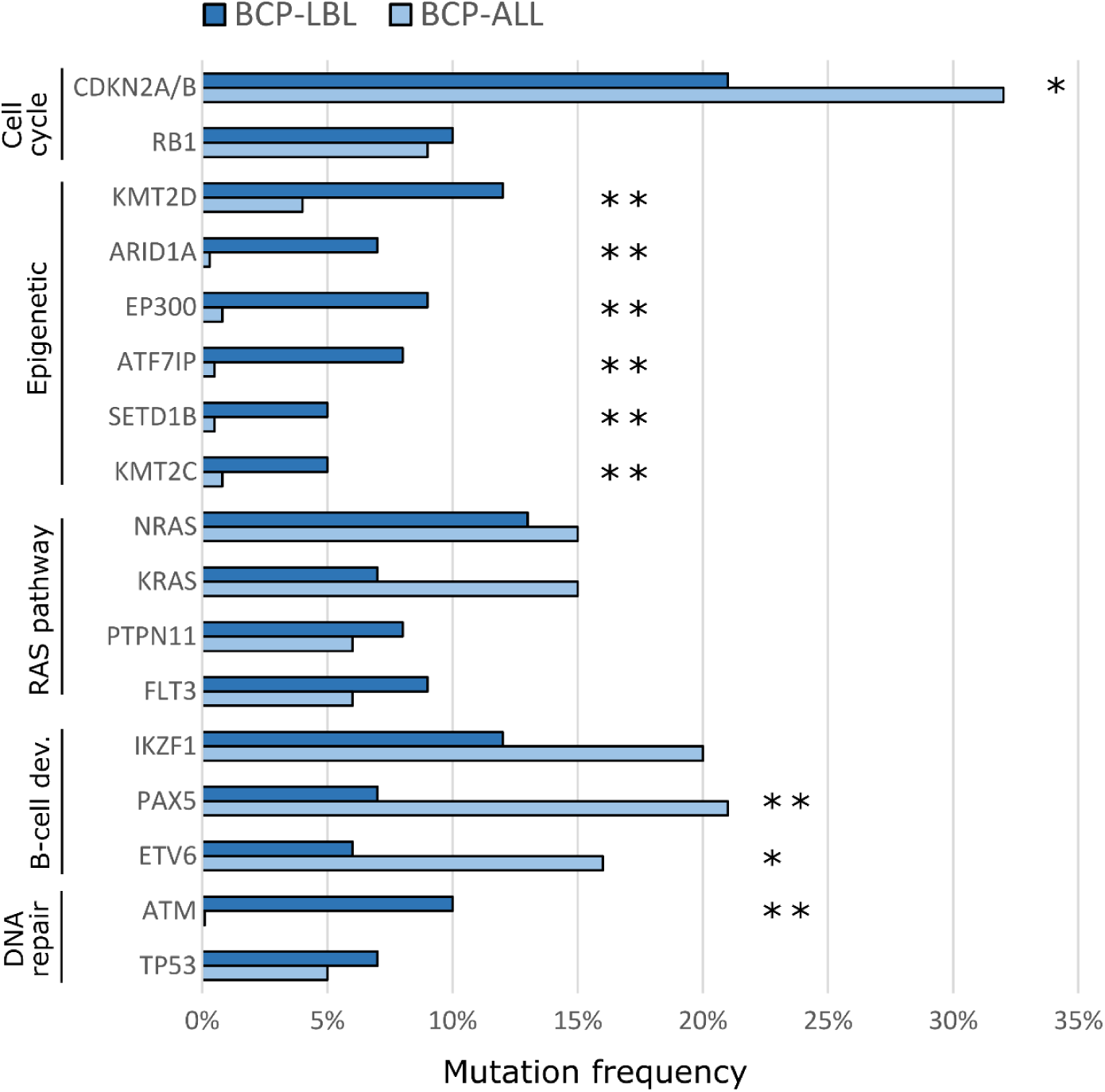
Comparison of the frequencies of gene aberrations and genetic subtypes between BCP-LBL and BCP-ALL. Most frequently mutated genes in current BCP-LBL cohort were compared with mutational frequencies of these genes in 608 BCP-ALL patients from Brady et al. (2022). Asterisk indicates statistical significant difference between mutational frequency in BCP-LBL versus BCP-ALL (* < 0.05; ** < 0.005)

### Integrating copy number and gene expression to define BCP-LBL subtypes

Given the overlap in cell of origin between BCP-LBL and BCP-ALL, we evaluated whether the subtype classification of BCP-ALL^1^ could be applied to BCP-LBL patients. Subtypes were defined for 87 samples by integrating DNA CNAs, SNVs and gene fusions. Additionally, we used gene expression to predict subtypes for 80 BCP-LBL cases by RNAseq data using the machine learning-based ALLCatchR classifier^15^, which was developed on BCP-ALL. Combining these approaches, we classified 81 BCP-LBL patients to one of 17 subtypes leaving only 6 (7%) patients without subtype-defining lesions as ‘BCP-LBL not otherwise specified’ (NOS) (S**upplemental Table 3**). Ten patients with incomplete data due to technical limitations of the material could not be assigned to a subtype. The successfully classified BCP-LBL could be lumped into groups based on aneuploidies (40%), TF-driven aberrations (46%) and kinase/cytokine receptor-driven aberrations (7%). These rates were comparable with data from BCP-ALL in which aneuploidy-based account for 34%, TF-driven for 44% and kinase/cytokine-driven for 12% of subtypes^7^ (**Figure 6A**). Using reference subtypes based on aneuploidies, gene fusions and SNVs, we evaluated the overall prediction accuracy of ALLCatchR applied to BCP-LBL as 87.5% (**Figure 6B**). The specificities for all subtypes were above 90%, suggesting that false-positive predictions were rare. For several BCP-LBL subtypes, including iAMP21 and near haploidy, the classifier had a lower sensitivity (**Figure 6C**), which was in line with the sensitivities reported for BCP-ALL^15^. Among the aneuploidy-driven subtypes, the HeH subtype was the one predicted by the ALLCatchR classifier with the highest accuracy (81%). The iAMP21 BCP-LBL subtype was predicted with 70% accuracy, with the remaining cases classified as Ph-like. (**Figure 6B**). More than 80% of the TF-driven subtypes and all the kinase/cytokine receptor-driven subtypes, except for one, were classified correctly by the ALLCatchR classifier. Finally, for each of the identified subtype, ALLCatchR could predict a cell of origin based on enrichment scores for the seven B-cell differentiation stages (**Supplemental Figure 3**).

**Figure 6:**
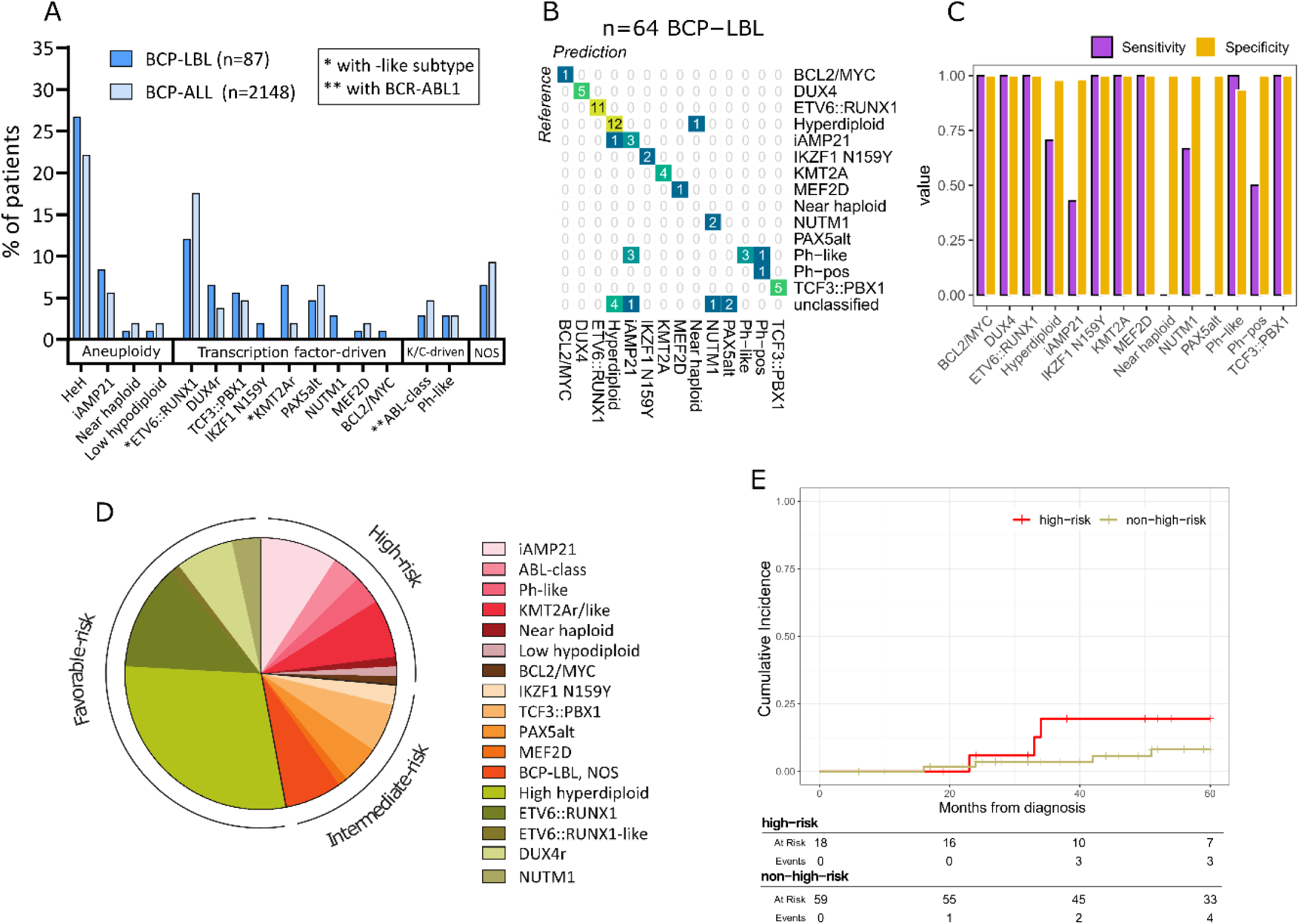
Overview of subtypes and genetic-risk group in BCP-LBL. (A) Comparison of frequencies of subtypes in BCP-LBL compared to published data from Brady et al. (2022) showing a largely similar distribution. (B) Confusion matrix showing a high degree of concordance between the subtypes assigned based on aneuploidies, gene fusions and SNVs (reference, columns) and subtypes predicted using the ALLCatchR classifier (rows), that is solely based on gene expression data. (C) Bar graph showing for each of the subtype, the sensitivity and specificity of the ALLCatchR prediction. (D) Distribution of subtypes associated with risk for BCP-LBL based on known risk-associated subtypes in BCP-ALL. (E) Cumulative incidence of relapse of BCP-LBL cohort based on genetic-risk groups known from BCP-ALL, comparing high-risk versus non-high-risk genetic subtypes. Outcome data or follow-up data was missing for 20 patients.

### Comparison of subtypes in BCP-LBL and BCP-ALL Subtypes based on aneuploidies

The most prevalent subtype in the BCP-LBL cohort was HeH, observed in 25 out of 87 patients (29%). HeH is also the most common in pediatric BCP-ALL, occurring in approximately a quarter of the patients. ^7^ We observed RAS pathway mutations (*KRAS*, *NRAS*, *PTPN11*, *FLT3*) in 32% of the HeH patients. The iAMP21 subtype occurred in 8 out of 87 BCP-LBL patients (9%), comparable to the reported prevalence in BCP-ALL (6%). A gain of chromosome 21 was also found in subtypes other than HeH and iAMP21, making it the most frequent copy number-altered chromosome in BCP-LBL (48%). The low hypodiploid subtype was detected in 1 out of 87 patients. Lastly, the near haploid subtype was observed in 1 out of 87 patients in our cohort. These frequencies are comparable to those in BCP-ALL (**Figure 6A and Supplemental Table 6**).

### TF-driven subtypes

The most common TF-driven subtype in our BCP-LBL cohort was *ETV6*::*RUNX1*, detected in 11 out of 87 cases (13%). *ETV6*::*RUNX1* fusions are detected in pediatric BCP-ALL with a frequency of 19% in the BCP-ALL comparison cohort and a frequency reaching 25% in literature^6^. Although not significant (**Supplemental Table 6**), the percentage of BCP-LBL patients with the *ETV6*::*RUNX1* subtype seems to be lower than in BCP-ALL. The median age of BCP-LBL patients in the *ETV6*::*RUNX1* subtype was 6 years. Additional genetic alterations have been found to be very heterogeneous in patients in the *ETV6*::*RUNX1* subtype, but approximately 70% of the BCP-ALL patients have *ETV6* deletions. We did not detect *ETV6* deletions in our *ETV6*::*RUNX1* BCP-LBL cases, which could be due to the limited ability to detect focal deletions in WES data from FFPE samples.

Other TF-driven BCP-LBL subtypes were detected, which also occur in BCP-ALL: *DUX4* rearrangement (6 patients, 7%), *TCF3*::*PBX1* (5 patients, 6%), *PAX5* alteration (4 patients, 5%), *KMT2A* rearrangement (4 patients, 5%), *NUTM1* rearrangement (3 patients, 3%), *IKZF1* N159Y (2 patients, 2%), *MEF2D* rearrangement (1 patient, 1%), and *MYC* rearrangement (1 patient, 1%). *KMT2A*-rearranged BCP-LBL was as infrequent as in BCP-ALL in this cohort. The difference to our previous publication, in which we reported 19% (9/48) *KMT2A*-rearranged BCP-LBL^9^, is most likely due to the low number of cases with cutaneous biopsies in the current study, and in the different techniques (FISH vs RNAseq) used to identify the gene fusions in the two studies. We also detected two *KMT2A*-like cases, that were classified as *KMT2A*-rearranged by ALLCatchR and characterized by *MLLT10* rearrangements and increased *HOXA9*/*HOXA10* expression.

### Kinase/cytokine receptor-driven subtypes

The last major group of BCP-ALL is characterized by tyrosine kinase/cytokine receptor activation, resulting in constitutive kinase activity. In our BCP-LBL cohort, we observed *ABL*-class tyrosine kinase fusions *BCR*::*ABL1* (n=2) and *ETV6*::*ABL1* (n=1), JAK-class tyrosine kinase fusion *PAX5*::*JAK2* (n=1) and cytokine receptor fusion *P2RY8*::*CRLF2* (n=1). Furthermore, one BCP-LBL sample out of 87 showed overexpression of *CRLF2* without a chimeric transcript, possibly caused by a rearrangement of *CRLF2* to the IGH enhance. Together, kinase/cytokine receptor-driven subtypes constituted 6% of the BCP-LBL cohort, which was comparable to BCP-ALL (8%).

For all of the identified subtypes, the B-lymphopoiesis trajectories predicted by the ALLCatchR classifier were strikingly similar to those predicted on a pediatric BCP-ALL cohort (**Supplemental Figure 3**).

In BCP-ALL, genetic risk groups have been defined with high-risk subtypes: iAMP21, ABL-class, *KMT2A*-rearranged, *KMT2A*-like, near haploid, low hypodiploid, *MYC*-rearranged; intermediate-risk subtypes: *IKZF1* N159Y, *TCF3*::*PBX1*, *PAX5*-altered, *MEF2D*-rearranged, BCP-ALL, NOS; favorable-risk subtypes: *DUX4*-rearranged, HeH, *ETV6*::*RUNX1*, *NUTM1*-rearranged (**Figure 6D**). Applying these categories to BCP-LBL, we identified high-risk genetics in 24% of BCP-LBL cases. More relapses were observed in the group with high-risk genetics compared to the group without high-risk genetics but the cumulative incidence of relapse did not reach statistical significance (p=0.2) (20%; 95% confidence interval [CI], 4.4 to 43 vs.8.1%; 95% CI, 2.5 to 18) (**Figure 6E**). There was no association between stage of disease and genetic risk groups, as the distribution of stage I/II and stage III/IV was similar in high-risk and non-high-risk groups (**Supplemental Figure 2**).

## Discussion

This is the first study to date describing the genomic and transcriptomic landscape of pediatric BCP-LBL. Tissue material from BCP-LBL patients is usually limited, hampering comprehensive molecular subtyping. However, we showed that the combination of WES and RNAseq is able to detect genetic subtypes, even in FFPE material. Our approach provides insight into the biology of BCP-LBL, its relation to BCP-ALL and may also provide a basis for future diagnosis and risk stratification.

BCP-LBL is a malignancy of immature precursor B-cells, as demonstrated by analysis of IG and TR rearrangements, without any obvious difference to BCP-ALL regarding the maturation stage of the cell of origin. Although some studies suggested that structural abnormalities such as translocations are more prevalent in leukemia, whereas numerical chromosome aberrations are more prevalent in lymphoma ^10,29^, this does not seem to apply to BCP neoplasms. E.g., the frequency of aneuploidy-associated subtypes in BCP-LBL were not significantly different to BCP-ALL in our study. We found a broad overlap of somatic mutations between BCP-LBL and BCP-ALL. Nevertheless, for some of the mutated genes, frequencies differed significantly between leukemia and lymphoma. Whereas alterations in genes encoding TFs of B-cell development, such as *PAX5* and *ETV6* were significantly more frequent in BCP-ALL, genes encoding epigenetic regulators (*ARID1A*, *EP300*, *KMT2D*) were significantly more often mutated in BCP-LBL. This finding is of potential clinical relevance given that changes in the epigenome may be linked to drug resistance^28^. We demonstrate that virtually all molecular subtypes identified in BCP-ALL were also present in BCP-LBL. We also show that a prediction tool trained on BCP-ALL gene expression data reliably identified subtypes in BCP-LBL, further stressing the genetic similarity between BCP-LBL and BCP-ALL. By comparing genetic risk-groups based on BCP-ALL, we show that BCP-LBL patients with high-risk genetics have a higher cumulative incidence of relapse, a finding that did not reach statistical significance, as the number of cases and events were too small for definite conclusions. High-risk molecular subtypes occurred independent of disease stage but single cases of stage I/II disease showed high risk genetics. Thus, the clinical relevance and interdependence of stage - which is currently used for stratification of therapy - and high-risk by molecular profiling is still uncertain and requires larger cohorts to be analyzed.

In addition to finding fusions known from BCP-ALL, our study also led to the identification of novel fusions involving genes both known and not known in BCP-ALL like hitherto unknown fusion partners of *TCF3* and *PAX5* (*GLIPR2* and *BRD3*, respectively). Another novel fusion involved the Mitochondrial Translation Optimization 1 (*MTO1*) and the membrane localization domain of Junctophilin 3 (*JPH3*). One might speculate on a possible effect of this alteration on metabolism and protein translation but further studies are needed to understand its biological significance.

There are several limitations of our study. First, germline DNA for most historic samples was not available and we thus restricted the analysis to genes known to be drivers in BCP-ALL complemented with lymphoma genes and DNA repair genes to filter the BCP-LBL WES data. Thus, we cannot exclude that genes not previously reported in BCP-ALL may be mutated in BCL-LBL. Future analyses need to address this question, especially in BCP-LBL in which no driver mutations known from BCP-ALL have been identified. Additionally, we found that a significant number of our BCP-LBL cases carried mutations in DNA repair genes. Future studies on paired tumor and normal DNA are needed to show whether germline mutations in DNA repair genes may play a role in BCP-LBL, similarly to what observed in BCP-ALL^30^. Second, despite the absence of a bias in respect to clinical parameters compared to previously published cohorts, our study may suffer from a selection bias due to the localization of the biopsies. A smaller cohort of BCP-LBL previously analyzed by FISH, included 17% cases with cutaneous biopsies showing a frequency of *KMT2A* breakpoints of 19%^9^, while in our current cohort with only 8% cutaneous biopsies, the frequency of *KMT2A* fusions was 5%. Understandably, cutaneous biopsies of pediatric patients are small and may often not qualify for molecular profiling as conducted in the current study. Similarly, the frequency of BCP-LBL with ABL-class fusions needs to be finally determined in future studies, using RNAseq or break-apart FISH for the ABL class kinase genes *ABL1*, *ABL2*, *PDGFRB* and *CSF1R*. Whether these patients benefit from including imatinib into the treatment protocols needs to be determined. Nevertheless, our study shows that a molecular subtyping of BCP-LBL in analogy to BCL-ALL may be helpful for individualized treatment decisions.

In summary, by carrying out a detailed molecular analysis of a large cohort of patient samples, we defined the mutational and transcriptional landscape of BCP-LBL. Our study revealed more similarities than differences between BCP-LBL and BCL-ALL reinforcing the concept of two manifestations of the same entity. In addition, the striking biological similarities between BCP-LBL and BCP-ALL challenges the concept of classification as leukemia or lymphoma based on the extent of bone marrow infiltration. Moreover, our study provides a blueprint how to implement molecular analysis of BCP-LBL and thus presents the basis for further studies aimed at defining new, genetics-based treatment protocols, in which BCP-LBL patients may also benefit from new BCP-ALL treatment strategies.

## Acknowledgments

We thank Dana Germer, Charlotte Botz von Drathen, Reina Zuhlke-Jenisch, and Lorena Valles Uriarte for excellent technical support. We thank the Princess Máxima Center Biobank and the Dutch Pathology register PALGA for access to material. We thank the University Medical Center Utrecht Pathology Department and the Princess Máxima Center Diagnostic Laboratory for excellent technical support. We thank Jules Meijerink for his role in awarding the grant.

This work was supported by Stichting Kinderen Kankervrij (KiKa) grant number 393, the DFG-funded clinical research unit CATCH-ALL (https://www.catchall-kfo5010.com/), the KinderKrebs Initiative Buchholz, Holm-Seppensen (KKI), The Ferenc Foundation, and a grant from the family Onderwater. BB is supported by the Deutsche Kinderkrebsstiftung to conduct the NHL-BFM Registry 2012.

We are indebted to patients, parents and treating physicians for their contributions.

## Authorship Contributions

M.L.B., J.M.B., J.L.C.L., A.B., I.I. and W.K. designed the study;

L.A.K., M.K., M.M., M.P.H., and R.S.B. performed and analyzed sequencing experiments;

E.K., I.I., M.M.K., T.B., M.B., R.P.K., and J.M.B. analyzed and interpreted molecular and clinical data;

M.S.V. and W.K. analyzed and interpreted histopathology data;

N.G., A.A., J.J.V.A., R.S., E.M., A.B., and J.L.C.L. collected patient samples;

K.S., L.A.P, G.C. and B.B. provided patient data;

B.B., M.B., R.K., C.D.B, T.B., and L.A.P. provided important advice;

E.K., I.I., J.M.B., and W.K. wrote the manuscript; and all other authors reviewed and approved the final version of the manuscript.

## References

1. Alaggio R, Amador C, Anagnostopoulos I, et al. The 5th edition of the World Health Organization Classification of Haematolymphoid Tumours: Lymphoid Neoplasms. Leukemia 2022;36:1720–48.

2. Kroeze E, Arias Padilla L, Bakker M, et al. Pediatric Precursor B-Cell Lymphoblastic Malignancies: From Extramedullary to Medullary Involvement. Cancers 2022;14:3895.

3. Hunger SP, Mullighan CG. Acute Lymphoblastic Leukemia in Children. N Engl J Med 2015;373:1541–52.

4. Linabery AM, Ross JA. Trends in childhood cancer incidence in the U.S. (1992-2004). Cancer 2008;112:416–32.

5. Reedijk AMJ, Coebergh JWW, de Groot-Kruseman HA, et al. Progress against childhood and adolescent acute lymphoblastic leukaemia in the Netherlands, 1990-2015. Leukemia 2020.

6. Schwab C, Harrison CJ. Advances in B-cell Precursor Acute Lymphoblastic Leukemia Genomics. Hemasphere 2018;2:e53.

7. Brady SW, Roberts KG, Gu Z, et al. The genomic landscape of pediatric acute lymphoblastic leukemia. Nat Genet 2022;54:1376–89.

8. Schraders M, van Reijmersdal SV, Kamping EJ, et al. High-resolution genomic profiling of pediatric lymphoblastic lymphomas reveals subtle differences with pediatric acute lymphoblastic leukemias in the B-lineage. Cancer Genet Cytogenet 2009;191:27–33.

9. Sharma R, Klairmont MM, Holland AC, et al. Integrative genomic analysis of B-lymphoblastic lymphoma with intrachromosomal amplification of chromosome 21. Pediatr Blood Cancer 2020:e28357.

10. Knez V, Bao L, Carstens B, Liang X. Analysis of clinicopathological and cytogenetic differences between B-lymphoblastic lymphoma and B-lymphoblastic leukemia in childhood. Leuk Lymphoma 2020:1–7.

11. Kubota-Tanaka M, Osumi T, Miura S, et al. B-lymphoblastic lymphoma with TCF3-PBX1 fusion gene. Haematologica 2019;104:e35–e7.

12. Au-Yeung RKH, Padilla LA, Zimmermann M, et al. Frequency and prognostic implications of KMT2A rearrangements in children with precursor B-cell lymphoma. Leukemia 2023;37:488–91.

13. Stewart JP, Gazdova J, Darzentas N, et al. Validation of the EuroClonality-NGS DNA capture panel as an integrated genomic tool for lymphoproliferative disorders. Blood Adv 2021;5:3188–98.

14. DePristo MA, Banks E, Poplin R, et al. A framework for variation discovery and genotyping using next-generation DNA sequencing data. Nat Genet 2011;43:491–8.

15. Wood RD, Mitchell M, Sgouros J, Lindahl T. Human DNA repair genes. Science 2001;291:1284–9.

16. Vaque JP, Martinez N, Batlle-Lopez A, et al. B-cell lymphoma mutations: improving diagnostics and enabling targeted therapies. Haematologica 2014;99:222–31.

17. Van der Auwera GA, Carneiro MO, Hartl C, et al. From FastQ data to high confidence variant calls: the Genome Analysis Toolkit best practices pipeline. Curr Protoc Bioinformatics 2013;43:11 0 1-0 33.

18. Dobin A, Davis CA, Schlesinger F, et al. STAR: ultrafast universal RNA-seq aligner. Bioinformatics 2013;29:15–21.

19. Uhrig S, Ellermann J, Walther T, et al. Accurate and efficient detection of gene fusions from RNA sequencing data. Genome Res 2021;31:448–60.

20. Beder T, Hansen BT, Hartmann AM, et al. The Gene Expression Classifier ALLCatchR Identifies B-cell Precursor ALL Subtypes and Underlying Developmental Trajectories Across Age. Hemasphere 2023;7:e939.

21. Burkhardt B, Zimmermann M, Oschlies I, et al. The impact of age and gender on biology, clinical features and treatment outcome of non-Hodgkin lymphoma in childhood and adolescence. Br J Haematol 2005;131:39–49.

22. Landmann E, Burkhardt B, Zimmermann M, et al. Results and conclusions of the European Intergroup EURO-LB02 trial in children and adolescents with lymphoblastic lymphoma. Haematologica 2017;102:2086–96.

23. Campo E, Jaffe ES, Cook JR, et al. The International Consensus Classification of Mature Lymphoid Neoplasms: a report from the Clinical Advisory Committee. Blood 2022;140:1229–53.

24. Oschlies I, Burkhardt B, Chassagne-Clement C, et al. Diagnosis and immunophenotype of 188 pediatric lymphoblastic lymphomas treated within a randomized prospective trial: experiences and preliminary recommendations from the European childhood lymphoma pathology panel. Am J Surg Pathol 2011;35:836–44.

25. Krangel MS. Gene segment selection in V(D)J recombination: accessibility and beyond. Nat Immunol 2003;4:624–30.

26. Kotrova M, Darzentas N, Pott C, Baldus CD, Bruggemann M. Immune Gene Rearrangements: Unique Signatures for Tracing Physiological Lymphocytes and Leukemic Cells. Genes (Basel) 2021;12.

27. Szczepanski T, Beishuizen A, Pongers-Willemse MJ, et al. Cross-lineage T cell receptor gene rearrangements occur in more than ninety percent of childhood precursor-B acute lymphoblastic leukemias: alternative PCR targets for detection of minimal residual disease. Leukemia 1999;13:196–205.

28. Saint Fleur-Lominy S, Evensen NA, Bhatla T, et al. Evolution of the Epigenetic Landscape in Childhood B Acute Lymphoblastic Leukemia and Its Role in Drug Resistance. Cancer Res 2020;80:5189–202.

29. Maitra A, McKenna RW, Weinberg AG, Schneider NR, Kroft SH. Precursor B-cell lymphoblastic lymphoma. A study of nine cases lacking blood and bone marrow involvement and review of the literature. Am J Clin Pathol 2001;115:868–75.

30. Elitzur S, Shiloh R, Loeffen J, et al. ATM Germline Pathogenic Variants Affect Treatment Outcomes in Children with Acute Lymphoblastic Leukemia/Lymphoma and Ataxia Telangiectasia. Blood 2023;142:520.

