## Supplemental Figures for "Mutational and transcriptional landscape of pediatric B-cell precursor lymphoblastic lymphoma"

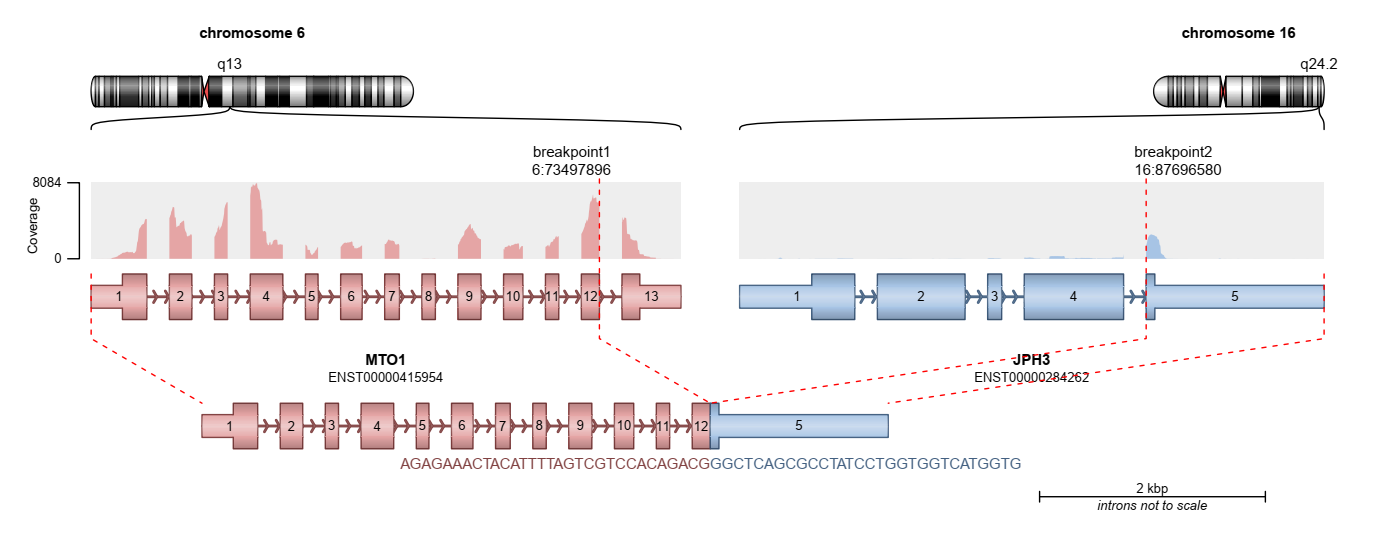

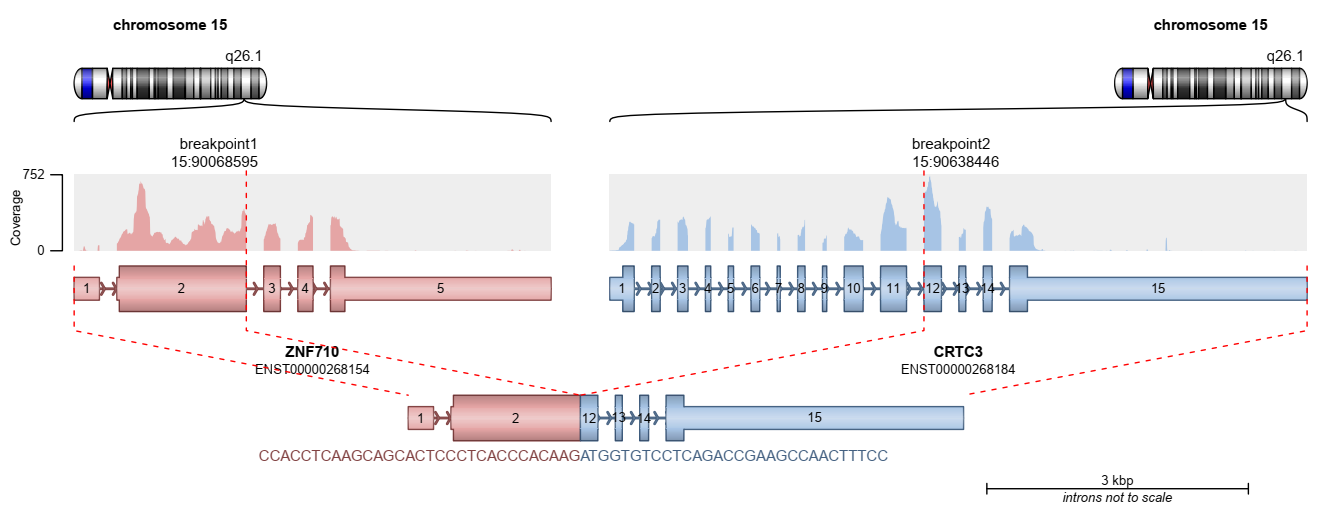

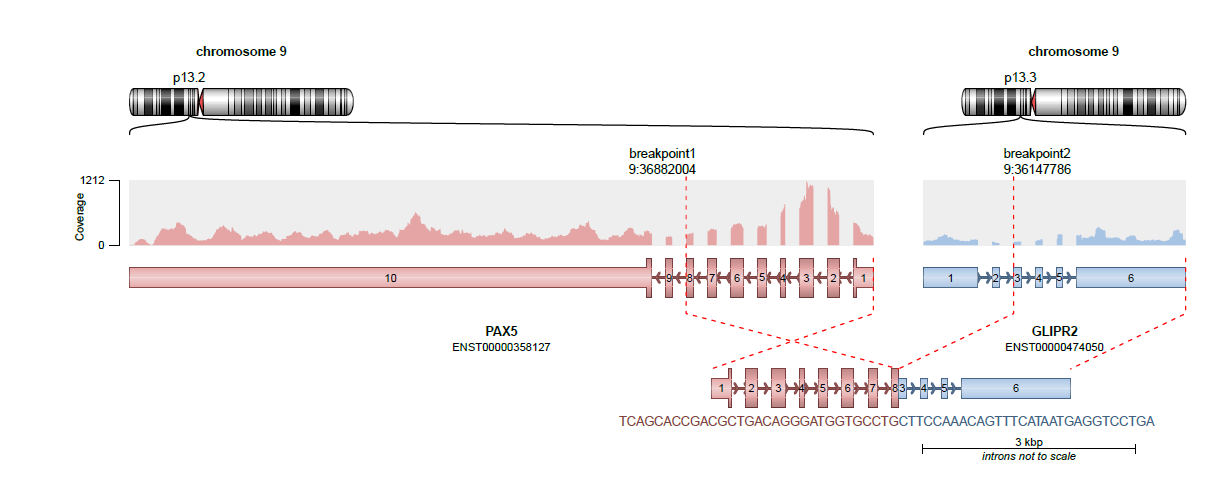

**Supplemental Figure 1**: Arriba visualization of the newly identified fusion between the MTO1 and JPH3 genes, between the ZNF10 and CRTC3 genes and between the PAX5 and GLIPR2 genes.

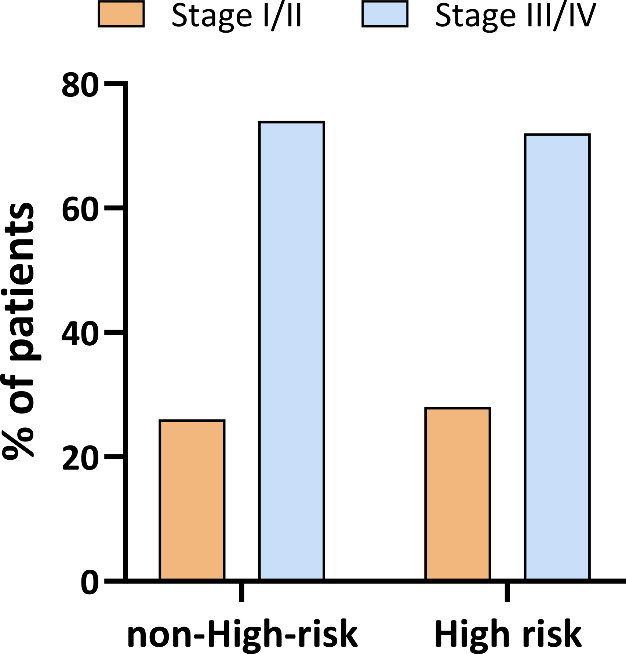

**Supplemental Figure 2**: Association between risk-associated genetics and stage

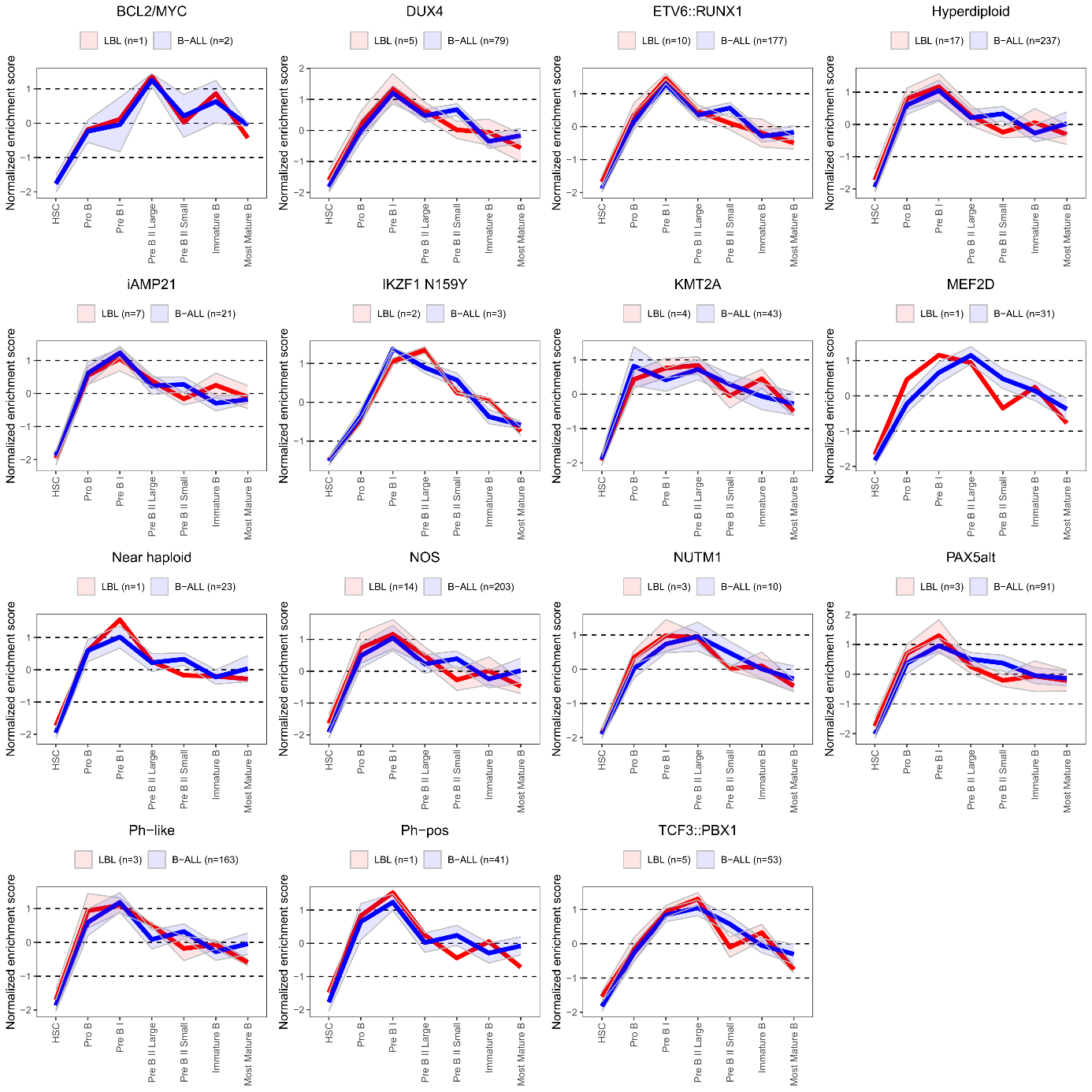

**Supplemental Figure 3**: Developmental trajectories of pediatric BCP-LBL in comparison to BCP-ALL. Enrichment scores (similarity value) were derived using the ALLCatchR predictor, based on gene set enrichment analysis of gene sets specific for the B cell stages. The figure shows the enrichment scores (y-axis) for the 7 B-cell stages (x-axis) comparing subtypes of pediatric BCP-LBL (red line) with pediatric BCP-ALL (blue line). Shade around lines indicate standard deviation.
