## Supplemental Methods for "Mutational and transcriptional landscape of pediatric B-cell precursor lymphoblastic lymphoma"

*Immunohistochemistry*

Immunophenotyping of samples was achieved using immunohistochemistry with antibodies directed against cluster of differentiation (CD)19 and/or CD79a, as B cell markers; CD3, as a general T cell marker; terminal deoxynucleotidyl transferase (TdT) as marker for precursor lymphoid neoplasms, and myeloperoxidase (MPO) as marker for the myeloid lineage. Immunohistochemistry was performed on an automated stainer using protocols recommended by the supplier as previously published^1^.

*Whole exome sequencing (WES)*

FFPE material from the Netherlands was collected from the Dutch Nationwide Pathology Databank (DNPD)^2^. Tumor percentage in FFPE biopsies was estimated by hematoxylin and eosin (HE) staining combined with pathology reports. Genomic DNA extraction from the samples processed in the Netherlands was achieved using Maxwell FFPE Plus DNA kit (Promega, Mannheim, Germany) and quantified using a Qubit 2.0 Fluorometer (Thermo Fisher Scientific, CA, USA) according to manufacturer’s instructions. DNA extracted from FFPE tissue was sheared by ultrasound with a Covaris S2 (Covaris Inc, MA, USA) into aimed lengths of 180-200 base pairs (bp). DNA from viably frozen material was enzymatically fragmented. Library preparation was conducted using KAPA Library Preparation (KAPA Biosystems, MA, USA) and HyperPlus Capture Exome Kit. Samples were sequenced on the Illumina Novaseq6000 (2x150bp paired-end). For samples collected in Germany, genomic DNA was extracted using the PureLink® Genomic DNA Kit (Thermo Fisher Scientific, CA, USA) and quantified using a Qubit 2.0 Fluorometer (Thermo Fisher Scientific, CA, USA). DNA quality was assessed by capillary electrophoresis (TapeStation, Agilent) and qPCR (Infinium HD FFPE QC Assay, Illumina, San Diego, CA, USA). Libraries for next generation sequencing (NGS) were prepared using the KAPAHyperPlus Kit (KAPA Biosystems, MA, USA) using a workflow optimized for FFPE DNA and captured with the xGen Exome Panel V2 (Integrated DNA Technology, USA) using the xGen Hybridization and Wash Kit (Integrated DNA Technology, USA). Pools of 16 libraries were sequenced on one lane of the Illumina Novaseq6000 (2x150bp paired-end). The aimed coverage for samples from both cohorts was approximately 200X.

*WES data analysis*

The reads were aligned to Hg38 using BWA, duplicate reads were marked with Samtools 1.9^3,4^, and somatic mutations were called using Mutect2 from GATK 4.2.0.0^5^. To merge the two datasets, the intersect of the xGen Exome Panel V2 and HyperPlus Capture Exome Kit was taken. Ensembl VEP v105 was used to annotate the detected mutations^6^. Data were filtered on GnomAD v2.1 population frequency (popMax) ≤0.01, a variant allele frequency (VAF) of ≥0.2, combined with an alternative allele count of ≥10. Mutations with a Combined Annotation Dependent Depletion (CADD) score >15 were considered to be potentially pathogenic and included for further analysis. Due to the absence of germline material, we focused the mutation analysis on a selection of 536 genes with known oncogenic potential. A list of genes used for variant analysis can be found in **supplemental Table 1.** Genes can be classified as BCP-ALL candidate genes^7^, genes with a known function in DNA repair or genes frequently mutated in lymphomas^8,9^. For visualization of mutations, an oncoplot was generated using the ComplexHeatmap package^10^. Mutations that passed filtering steps and that were found to occur in ≥5% of the samples were visualized. Copy number aberrations were called by implementing GATK v4.0.1.2 best practices^11^.

*RNA sequencing (RNAseq)*

RNAseq data for the samples processed in the Netherlands was obtained from the diagnostic laboratory of the Princess Máxima Center. RNA was extracted using Maxwell RSC RNA FFPE kit (Promega, Mannheim, Germany) according to manufacturer’s instructions and quantified using a Qubit 2.0 Fluorometer (Thermo Fisher Scientific, CA, USA). Total RNA was isolated using the AllPrep DNA/RNA/miRNA universal Kit (Qiagen) according to manufacturer’s instructions. RNA-seq libraries were generated using 300 ng RNA, with the KAPA RNA HyperPrep Kit with RiboErase (KAPA Biosystems, MA, USA). Libraries were sequenced on the Illumina Novaseq6000 (2x150bp paired-end). For samples collected in Germany, RNA was extracted using the ExpressArt FFPE Clear RNAready Kit (AmpTec, Hamburg, Germany), or the RecoverAll™ Total Nucleic Acid Isolation Kit for FFPE (Thermo Fisher Scientific, CA, USA) and quantified using a Qubit 2.0 Fluorometer (Thermo Fisher Scientific, CA, USA). RNA quality was assessed by capillary electrophoresis (TapeStation, Agilent). Only RNA samples with a DV200 (percentage of fragments of >200 nucleotides) higher than 30% were considered for sequencing. To improve the success rate of RNA-seq from low quality RNAs from FFPE, an exome capture RNA-seq approach was used^12^. Total RNA libraries were prepared with the xGen Broad-Range RNA Library Prep Kit (Integrated DNA Technology, USA). To select for sequences of coding genes, libraries were captured in pools of four with the xGen Exome Panel V2 (Integrated DNA Technology, USA) using the xGen Hybridization and Wash Kit (Integrated DNA Technology, USA).

*RNA-seq data analysis and subtype definition*

Pre-processing of the data from the Netherlands was done with standardized and in-house pipelines and guidelines^13^. In summary, STAR v2.7.2d was implemented for read assembly to GRCh38 (gencode version 31), STAR-Fusion v1.8.0 for gene fusion detection and Rsubread v1.6.4 for quantifying the reference transcriptome and generating count matrices. Samples that passed with 30-115 million unique reads were included for fusion analysis. Normalized gene expression (TPM) data was used to predict subtypes using the ALLCatchR classifier^14^. Subtypes were assigned using the following approach: first we assigned a subtype based on the analysis of aneuploidies, fusions and SNVs (column S, Supplementary Table S4). Next, we run the ALLCathR classifier and defined a classifier prediction based on a prediction confidence (columns T and U). A final subtype (column V) was assigned by integrating the information coming from aneuploidies, fusions, SNVs and ALLCatchR prediction. Samples with inconsistencies in the data coming from the different sources were defined as BCP-LBL not otherwise specified (NOS). Samples for which the lack of experimental data prevented subtype definition were labelled as “n.a.” (not assigned). For those cases with missing experimental data but assigned to a subtype with “high-confidence” by ALLCatchR^14^, the ALLCatchR prediction was used as final subtype.

*Analysis of IG and TR gene rearrangements*

IG and TR rearrangement analysis was performed using the EuroClonality-NDC Assay (Univ8 Genomics. Belfast, UK), according to the manufacturer. Briefly, 65 ng of each DNA library (the same used for whole exome sequencing) were pooled and captured with 4 µl of the EuroClonality-NDC Assay, using the KAPA HyperCapture Reagent Kit and the KAPA HyperCapture Bead Kit (KAPA Biosystems, MA, USA). The target-enriched pool was sequenced on an Illumina NextSeq 500/550 system using a 75 bp paired-end strategy performed on a 150-cycle NextSeq 500/550 Mid Output Kit (Illumina, San Diego, CA, USA). Data were analyzed using the ARResT/Interrogate bioinformatic platform (http://arrest.tools/interrogate-latest/).

*Comparison to BCP-ALL*

Frequencies of subtypes were compared with BCP-ALL patients aged 1-18 years (n=2148) from published data by Brady et al. (2022)^7^ (Childhood SR=52%, Childhood HR=42%, AYA=6%). Mutational data was compared with a subset of these patients for whom additional data on single nucleotide variants, insertions/deletions and structural variants was available (n=608). Statistical significance between frequency of mutations detected in BCP-LBL and BCP-ALL was calculated using the Chi-square test.

*Survival analysis*

A competing risk model was used to estimate the cumulative incidence of relapse from first diagnosis. Relapse was used as event and death in complete remission (1 case) and secondary malignancy (1 case) were used as competing events. The Gray’s test was used to compare the cumulative incidence of relapse between high-risk and non-high-risk genetics.

**References**

1.	Au-Yeung RKH, Padilla LA, Zimmermann M, et al. Frequency and prognostic implications of KMT2A rearrangements in children with precursor B-cell lymphoma. Leukemia 2023;37(2):488-491. DOI: 10.1038/s41375-022-01757-0.

2.	Casparie M, Tiebosch AT, Burger G, et al. Pathology databanking and biobanking in The Netherlands, a central role for PALGA, the nationwide histopathology and cytopathology data network and archive. Cell Oncol 2007;29(1):19-24. DOI: 10.1155/2007/971816.

3.	Li H, Durbin R. Fast and accurate short read alignment with Burrows-Wheeler transform. Bioinformatics 2009;25(14):1754-60. DOI: 10.1093/bioinformatics/btp324.

4.	Danecek P, Bonfield JK, Liddle J, et al. Twelve years of SAMtools and BCFtools. Gigascience 2021;10(2). DOI: 10.1093/gigascience/giab008.

5.	DePristo MA, Banks E, Poplin R, et al. A framework for variation discovery and genotyping using next-generation DNA sequencing data. Nat Genet 2011;43(5):491-8. DOI: 10.1038/ng.806.

6.	McLaren W, Gil L, Hunt SE, et al. The Ensembl Variant Effect Predictor. Genome Biol 2016;17(1):122. DOI: 10.1186/s13059-016-0974-4.

7.	Brady SW, Roberts KG, Gu Z, et al. The genomic landscape of pediatric acute lymphoblastic leukemia. Nat Genet 2022;54(9):1376-1389. DOI: 10.1038/s41588-022-01159-z.

8.	Wood RD, Mitchell M, Sgouros J, Lindahl T. Human DNA repair genes. Science 2001;291(5507):1284-9. DOI: 10.1126/science.1056154.

9.	Vaque JP, Martinez N, Batlle-Lopez A, et al. B-cell lymphoma mutations: improving diagnostics and enabling targeted therapies. Haematologica 2014;99(2):222-31. DOI: 10.3324/haematol.2013.096248.

10.	Gu Z, Eils R, Schlesner M. Complex heatmaps reveal patterns and correlations in multidimensional genomic data. Bioinformatics 2016;32(18):2847-9. DOI: 10.1093/bioinformatics/btw313.

11.	Van der Auwera GA, Carneiro MO, Hartl C, et al. From FastQ data to high confidence variant calls: the Genome Analysis Toolkit best practices pipeline. Curr Protoc Bioinformatics 2013;43(1110):11 10 1-11 10 33. DOI: 10.1002/0471250953.bi1110s43.

12.	Cieslik M, Chugh R, Wu YM, et al. The use of exome capture RNA-seq for highly degraded RNA with application to clinical cancer sequencing. Genome Res 2015;25(9):1372-81. DOI: 10.1101/gr.189621.115.

13.	Hehir-Kwa JY, Koudijs MJ, Verwiel ETP, et al. Improved Gene Fusion Detection in Childhood Cancer Diagnostics Using RNA Sequencing. JCO Precis Oncol 2022;6:e2000504. DOI: 10.1200/PO.20.00504.

14.	Beder T, Hansen BT, Hartmann AM, et al. The Gene Expression Classifier ALLCatchR Identifies B-cell Precursor ALL Subtypes and Underlying Developmental Trajectories Across Age. Hemasphere 2023;7(9):e939. DOI: 10.1097/HS9.0000000000000939.
